# Larger viral genome size facilitates emergence of zoonotic diseases

**DOI:** 10.1101/2020.03.10.986109

**Authors:** Richard E. Grewelle

## Abstract

Emergence of new viral diseases is linked to mutation or recombination events. The likelihood of cross-species transmission is related to phenotypic plasticity of a virus and its capacity to produce genetically variable progeny. Herein a model is described connecting the production of genetic variability with increasing genome size. Comparing all known zoonotic viral genome sizes to known non-zoonotic viral genome sizes demonstrates that zoonotic viruses have significantly larger genomes. These results support the notion that large viral genomes are important in producing new zoonotic disease, and suggest that genome size may be a useful surrogate in screening for potential zoonotic viruses.

## Introduction

Human population expansion continually redefines the interactions between humans and wildlife. Zoonotic diseases that emerge from these interactions account for over 60% of human infectious diseases [1]. Although this figure may represent over 2 billion new cases annually, zoonoses are characteristically underreported and misdiagnosed [2]. Because zoonoses affect humans and other vertebrates, they also generate substantial economic losses, as livestock death and human disability lead to loss of productivity [3]. For example, economic losses due to Ebola in 2014 were $2.2 billion [4]. Zoonotic outbreaks also present an increasingly pressing risk to human health due to higher densities of people with little to no immunity to novel pathogens. They are often unpredictable in occurrence and are caused by new species against which there are no developed vaccines, treatment, or public health protocol. This is particularly true for viruses that have great capacity to mutate, even during the course of a pandemic [1, 5]. This century has seen numerous pandemics caused by zoonotic viruses like Ebola, Nipah, Avian Influenza, and Coronavirus [5].

In late 2019, a novel Coronavirus (COVID-19) emerged and has since killed over 3000 people, making it the deadliest Coronavirus pandemic in recent history [6, 7]. The Wuhan province in China, where the virus emerged, effectively quarantined its populace, and many countries restricted travel of their own citizens to China. Although human to human transmission is driving current transmission, its emergence was facilitated by close human-animal contact, much like its predecessors SARS and MERS [8]. Members of the Coronavididae family are widely known to jump across host taxa to become infectious in new populations [9, 10]. Non-pathogenic varieties in a reservoir population can become pathogenic in new species. This phenomenon is well documented in many viral zoonoses, yet Coronaviruses pose exceptional risk to human populations due to the variety of potential reservoirs and the virulence of the emergent pandemics they cause [10]. The ability of Coronaviruses to shift to new species is due in part to the mutation rate they exhibit, but they are not exceptional among viruses in this way. They are exceptional, however, in their high frequency of recombination and in possessing the largest of the ssRNA viral genomes [11, 12]. These two characteristics provide ample opportunity for Coronaviruses to evolve new traits and establish themselves in new host species.

The capacity to produce genetic variability is a defining feature among viruses that are able to jump to new host species. Mutations and recombination events have been explored as a source of this variability. Both processes have been implicated in producing new viral strains or species responsible for observed pandemics [13]. The rates of mutation and recombination are known to vary across viral taxa and within viral genomes [14]. For example, RNA viruses characteristically mutate faster than DNA viruses due to polymerase infidelity, and hypermutable regions within some viral genomes have been demonstrated. Likewise, recombination processes vary considerably by viral type: reassortment is common in viruses with multi-segmented genomes, while homologous or non-homologous recombination is predominant in viruses with non-segmented genomes [14, 15].

A primary challenge for virologists and public health professionals is identifying characteristics of viruses that lead to higher likelihoods of new outbreaks [1]. The variety of mechanisms by which genetic variation can be produced makes finding clear patterns in zoonotic risk difficult [2, 13]. One yet unconsidered viral characteristic likely to facilitate cross-species infections is the size of the viral genome. Both the total number of mutations accumulated during replication and the probability of a recombination event increase with the size of a genome when replication mechanisms are conserved, though nucleotide substitutions from either process often lead to net reduction in fitness of progeny. The number of substitutions, thus potential for reduction in host-specific fitness, increases with genome size [16–18]. If genetic variability is instrumental in producing cross-species infections, genome size should be positively related to the likelihood of viral spillover events provided lethal substitutions do not substantially reduce the mean fitness of the viral population.

## Results

Cross-species transmission for each viral particle is assumed to be a function of the phenotypic plasticity of the viral genome and the genetic variability produced during replication [13, 15, 19]. The probability of a transmission event occurring can be represented as:

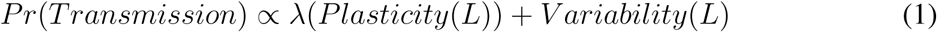

where *λ* ∈ [0, ∞) is the weighted contribution of plasticity over variability in promoting cross-species transmission and *L* is the length of the viral genome. Although a relationship between genome size and phenotypic plasticity is probable, little work has enlightened the nature of this relationship. What is clear is that transcription and gene regulation networks can become more complex with increasing number of nucleotides [20, 21]. Variability produced with each replication event can be more accurately approximated. Variability is produced via mutation and recombination (reassortment, non-homologous and homologous). Given per nucleotide rates of mutation (*µ*) and recombination (*R*), the accumulation of variability per genome should increase linearly as (*µ* + *R*(1 − *µ*))*L*. When *µ* and R are small, this variability can be expressed as (*µ* + *R*)*L*. Lethal mutagenesis and incompatible recombinants reduce viability of viral products. Viability converges to *e*^−*µγL*^ across mutational models [18] and is calculated as 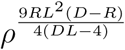 for recombination [17] (see supplement), where *γ* is the proportion of lethal mutations, *ρ* is the genome-specific tolerance to recombination events, and *D* ∈ [0, 1] is the sequence divergence between co-infecting viruses. Per nucleotide variability (*dV/dN*) produced with each replication event can be given as:

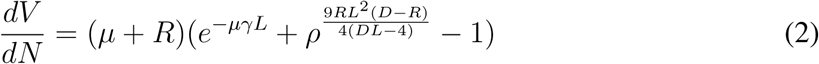

Per genome variability (*dV/dG*) can be given as:

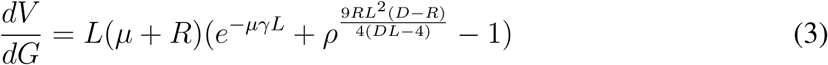

Assuming the number of polymerase enzymes scales with genome size [22, 23], a rate of production of variability can be derived.

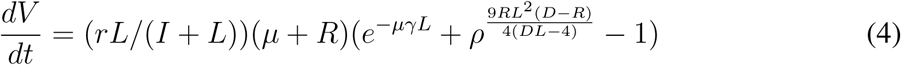

The efficiency of replication of each genome can be represented as *L/*(*I* + *L*), where *I* is the product of the time required for replication initiation and termination, and *r*, the rate of polymerase-mediated replication. Figure 1 displays equations 2–4 parameterized to represent positive sense ssRNA viruses. Per nucleotide variability production decreases with genome size due to reduced fitness with accumulated substitutions (Fig. 1A). However, genome-level variability production increases almost linearly with genome size, revealing that the rate of fitness reduction from the accumulation of deleterious substitutions does not outpace the increased variability associated with increase in genome size (Fig. 1B). Because more genomes can be replicated in a fixed amount of time for smaller genomes, the total variability produced in unit time gives indication of whether population-level variability increases with genome size. In fact, Fig. 1C shows that population-level variability is maximized within the known size range of ssRNA(+) viral genomes. Variability production per unit time continues to decline for genome sizes larger than 50,000 nt, though no known ssRNA viruses have genomes larger than 35,000 nt [12]. The maximum variability production per unit time is dependent on the efficiency of replication. Because Coronaviruses have much larger genomes than other ssRNA(+) viruses, they have evolved 3′ → 5′exoribonuclease repair and have lower mutation rates than other ssRNA(+) viruses [24]. Substituting this mutation rate in equation 4 produces a curve (Fig. S2) with a maximum shifted to the right, reflecting that production of genetic variability is maximized at a larger genome size for Coronaviruses than for other ssRNA(+) viruses. Considering these theoretical results, variability is greater per virion produced for viruses with larger genomes. Because population-level production of variability has been shown to improve viral survival via immune evasion [14], there is likely strong selection toward viral genome sizes close to the peak described by equation 4 (Fig. 1C). Even in the absence of phenotypic plasticity conferred by large genomes, viruses with large genomes have high potential to generate variability that promotes cross-species transmission events. Zoonotic viruses should, therefore, possess larger genomes than other viruses. To examine the relationship between viral genome size and likelihood of zoonosis, this study compiles a comprehensive list of all known viral zoonoses and leverages the entire NCBI database of viral genomes [12].

**Figure 1:**
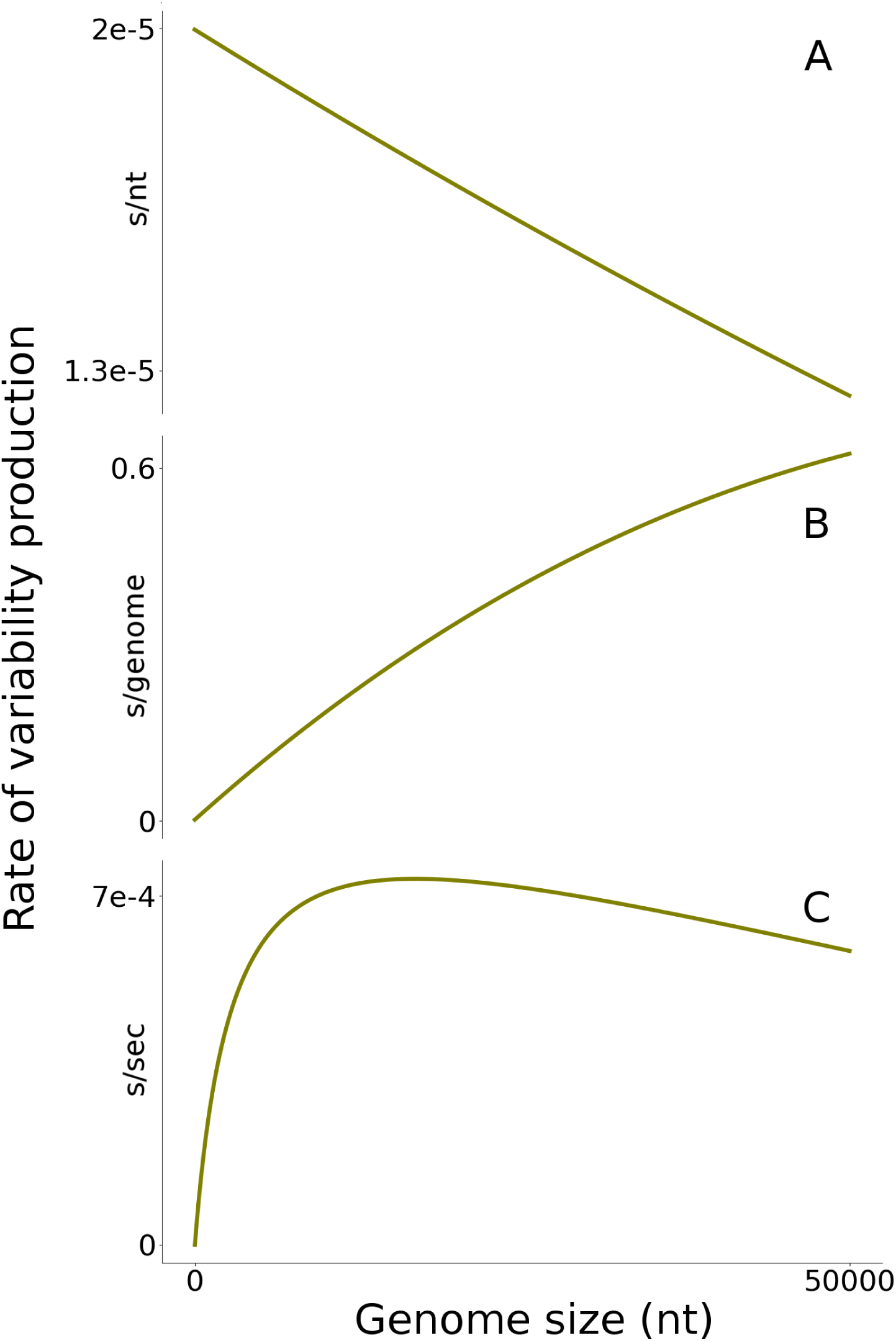
The rate at which genetic variability is produced per virion depends on genome size. The relationships are shown as substitutions per nucleotide (A), per genome (B), and per second (C) for a positive sense ssRNA virus.

Peer-reviewed literature from 1920 to 2019 was searched via the Web of Science electronic library for viral zoonoses. Search key words included “zoonotic”, “zoonoses”, “emerging disease”, “viral spillover”, “animal to human transmission”, and “animal reservoir virus”. After cross examination of all viruses yielded through this search, all confirmed zoonotic viruses transmitted from animals to humans in the past 100 years that also had a complete genome sequence were included in the analysis. This set encompasses 126 viral species from 40 genera in groups 1, 3, 4, 5, and 6 of the Baltimore classification system, grouped by genome structure. Genome sizes for all viruses were retrieved from the NCBI genome database or from peer-reviewed reports of fully sequenced genomes. The entire NCBI viral genome database of 9238 sequenced genomes was downloaded and filtered to remove incomplete, prokaryote, fungus, protist, and plant associated viral sequences. 2088 animal associated virus species in the five evaluated Baltimore groups remained after removal of zoonotic virus sequences. All species were categorized by genome structure according to the Baltimore classification system. One-tailed Welch’s T-tests for sample means were performed for each Baltimore group, comparing the genome sizes of zoonotic viruses with all other non-zoonotic viruses. Table 1 gives the results of these analyses, showing that zoonotic virus genomes are significantly larger than non-zoonotic virus genomes across viral groups. Resolution of this relationship is strong except for viral group 4, where the mean genome size for zoonotic viruses approaches being larger than the mean genome size for other animal viruses (at *α* = 0.05). The distributions of genome sizes are visualized in Fig. 2.

**Table 1:** Comparison of viral genome size between known zoonotic viruses and all other animal associated viruses. Comparisons are made within Baltimore groups classified according to genomic structure. Mean and standard errors for each group are reported.

| Group | Genome | Non-zoonotic Virus Genome Size | Zoonotic Virus Genome Size | P-value |
| --- | --- | --- | --- | --- |
| 1 | dsDNA | 75314 ± 3862 (n=665) | 165175 ± 12595 (n=8) | <0.0001 |
| 3 | dsRNA | 16065 ± 774 (n=106) | 21828 ± 2493 (n=4) | 0.05 |
| 4 | ssRNA(+) | 11205 ± 256 (n=662) | 12575 ± 920 (n=41) | 0.08 |
| 5 | ssRNA(-) | 10911 ± 184 (n=581) | 12964 ± 336 (n=64) | < 0.0001 |
| 6 | ssRNA-RT | 8664 ± 310 (n=74) | 10949 ± 637 (n=9) | 0.004 |

**Figure 2:**
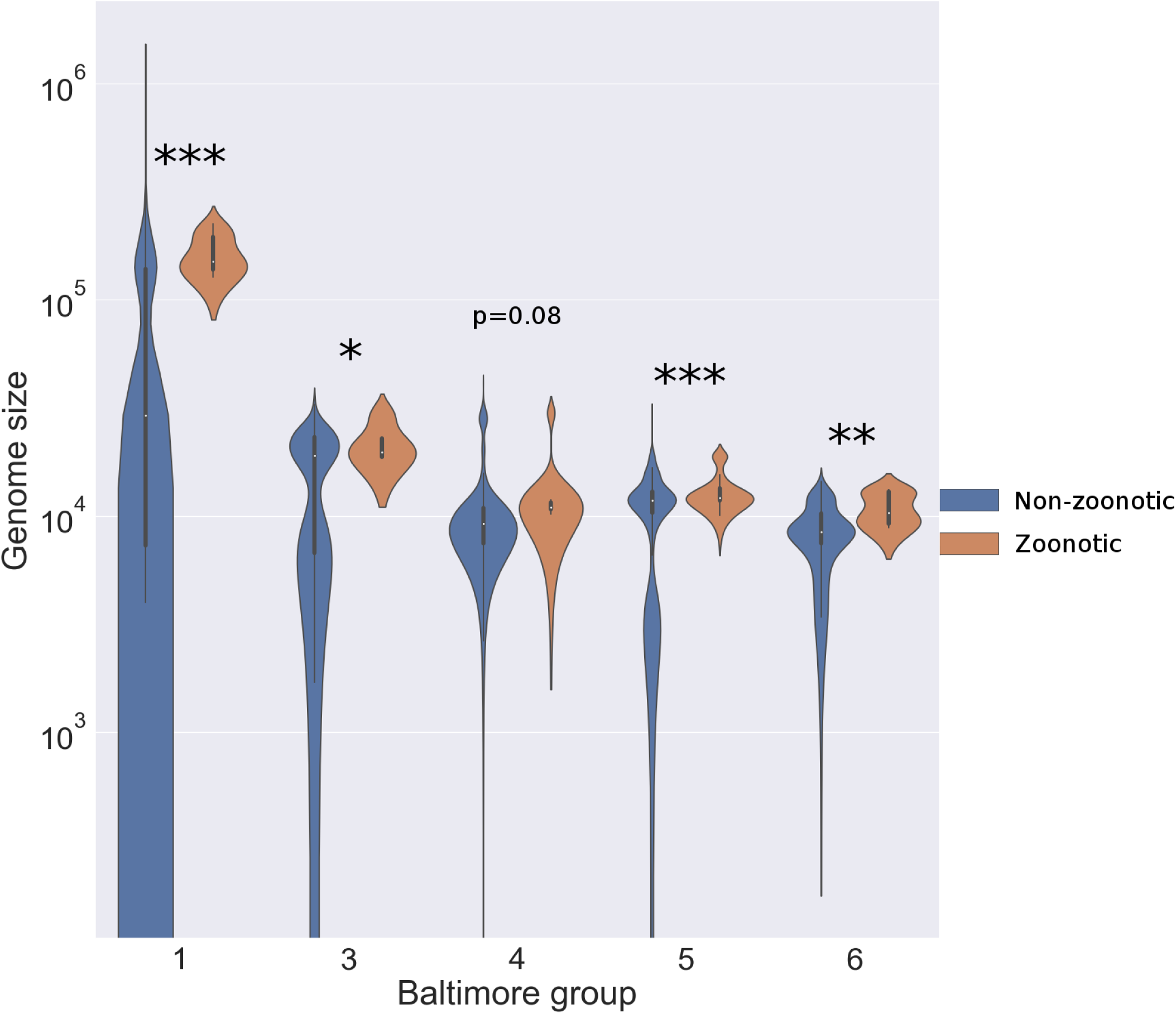
Violin plots comparing genome sizes between non-zoonotic animal viruses and zoonotic viruses. Zoonotic virus genome sizes inhabit the upper ranges of values in each group.

## Discussion

This finding strongly suggests that large genome size facilitates emergence of viral zoonoses. The signal is clear despite the large diversity of viruses within each Baltimore group and despite not accounting for viruses that share or jump between animal host species. Viruses with a wide host range likely share characteristics with zoonotic viruses, and based on these results, should have large genomes as well. For example, numerous species of Coronavirus are included in the non-zoonotic list (n=42) because they are not directly implicated in zoonoses, yet they are known to exhibit wide host ranges [9, 10]. Specifically, several bat Coronaviruses are in the non-zoonotic list, though researchers largely regard these viruses as the close relatives to SARS and MERS CoV [25]. This fact likely impaired the resolution of the relationship between genome size and cross-species transmission, resulting in the single non-significant p-value across evaluated groups. Large genomes are likely to facilitate zoonotic transmission via high mutational load per genome replication or high probability of recombination; both processes produce variability by which cross-species infection may occur. Genome sizes for some viral groups, especially dsDNA viruses, are tightly correlated with increasing number of genes (*R*^2^ = 0.91, Fig. S2). More coding regions within the genome could lead to greater plasticity and expanded host range, as reliance on host replication machinery is reduced [21, 26].

Among the viral groups there exists much variation in replication strategy, thus there is potential for differences in the predominant strategy for producing genetic variability leading to zoonotic transmission events [15]. Though mutation rate is 100 fold lower per nucleotide for dsDNA viruses than for RNA viruses, their much larger genomes enable a comparable likelihood of accumulating at least one mutation during replication. A negative correlation between mutation rate and genome size has been noted for a small set of viruses for which mutation rate has been directly measured [14]. Much of the correlation for RNA viruses results from measurement of bacteriophage mutation rate, and this interesting relationship should be further explored. Coronaviruses have evolved 3′→5′proofreading capabilities to accommodate mutagenesis associated with a large genome [24]. As recombination is generally associated with lower reduction in fitness compared to mutation, recombination may be the preferred mechanism to generate genetic variability for the viruses with the largest genomes. dsDNA viruses are known to have high per nucleotide rates of non-homologous recombination. Because of the large genome sizes of dsDNA viruses, the likelihood of at least one recombination event occurring during replication of the genome is high [15, 27]. The many coded genes of dsDNA viruses allow more easily incorporation of recombinant DNA without deleterious effects and may reduce host specificity through phenotypic plasticity [15, 17]. In contrast, Papillomaviruses are dsDNA viruses that are characteristically species-specific [28]. These viruses have small genomes with little plasticity to accommodate recombinant DNA or expand host range. dsRNA viruses are highly segmented, and the number of segments is highly correlated with the size of the genome (*R*^2^ = 0.84, Fig. S2), which allows for a high degree of recombination via reassortment of segments during a coinfection [15]. Gene expression is regulated by translation for ssRNA(+) viruses. This group includes Coronaviruses and is known for high levels of homologous recombination because of frequent template switching during discontinuous transcription. Having the largest RNA viral genomes, Coronaviruses can produce over 25% recombinant progeny with their high rates of recombination [11, 15]. ssRNA(-) viruses are not known to recombine with high frequency, so their genetic variation relies on high mutation rates alone [29]. ssRNA-RT viruses are known for maintaining high mutation rates characteristic of RNA viruses as well as high recombination rates for a subset of these viruses, including HIV. Pseudodiploidy in retroviruses can promote recombination due to the physical proximity of RNA strands in the virion as well as partial masking of deleterious mutations [30]. Large genomes are more likely to contain intergenic, non-coding regions that assist in gene regulation but also serve as hotspots for recombination and mutation due to weaker selection to conserve these regions [17].

Viral zoonoses can emerge as a result of large genetic changes associated with recombination or significant mutation–both more likely to occur in large genomes. Phenotypic plasticity is hypothesized to be positively related to genome size and may facilitate cross-species transmission, though this notion requires future exploration. The relationship between likelihood of zoonotic transmission and genome size has implications for predicting future zoonoses and for understanding general principles about viral life history. Host specificity may be considered a function of genome reduction. Guiding principles like the relationship between genome size and host range should be explored to broaden scientific understanding of the viral world and its role in ecosystems. Predicting future epidemics requires knowledge of potential zoonotic viruses. Far less than 1% of viruses have been discovered, and the potential host interactions of each viral species are even less well-characterized, making zoonotic risks difficult to estimate [31]. These data demonstrate that viral genome size is an effective indicator of zoonotic risk across viral genome subtypes. In an age of high-throughput genomics and rapid discovery of new viruses, characterization of recombination and mutation rates is far slower and more labor-intensive than genome sequencing, making genome size an appealing first step in zoonotic risk evaluation. Analysis of viral genome size and structure has the potential to provide deep insights in predicting future pandemics.

## Methods

### Derivation of Equation 2

The accumulation of substitutions in a genome occurs via mutation and recombination, with associated per nucleotide rates *µ* and *R*. These rates can be interpreted as the per nucleotide probability of a substitution occurring in a replication event. Assuming mutation and recombination are independent processes, the probability of a substitution (variability) at each nucleotide can be represented as:

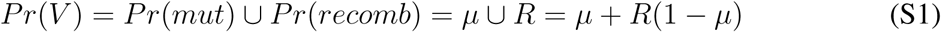

The probability that a substitution occurs for each nucleotide can be interpreted as a rate of substitution per replication event. Substitution events are assumed to be deleterious. Each event contributes to the reduction in fitness of the viral progeny, and these events accumulate in a genome, creating non-linear declines in fitness. Viral genome size is constrained by this reduction in fitness: larger genomes accumulate more substitutions and potential for fitness reduction is greater than for small genomes [14, 18]. Nucleotide-level fitness thus corresponds to genome-level fitness, and successful production of nucleotide variability is constrained by the fitness decay that occurs with increasing genome size. For purposes here, fitness will be synonymous with viability of each replicated genome. Setting the fitness of the parental virion or template genome at 1, fitness of progeny takes values between 0 and 1. Because a mutation and recombination event at a given nucleotide are mutually exclusive, the probability of each event occurring separately can be calculated as

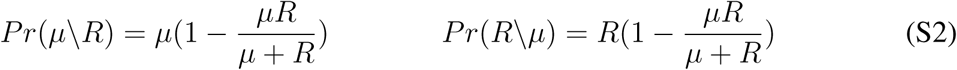

Across mutational models, fitness (infectivity) due to lethal mutagenesis is given as *ω*_*mut*_ = *e*^−*µγ*(*nt*)^ [18]. The fitness in the event of recombination is demonstrated to be 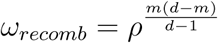 17, where *m* is the number of amino acid substitution events due to recombination and *d* is the maximum number of amino acid substitutions possible given the genetic dissimilarity of the coinfecting viral complement. Genome-level fitness for mutagenesis can be calculated as *ω*_*mut*_ = *e*^−*µγL*^. Because the probability of a random nucleotide substitution resulting in an amino acid substitution is 415*/*549 ≈ 3*/*4, the number of amino acid substitutions in a genome replication event is *m* = 3*RL/*4. Denoting *D* as proportional nucleotide dissimilarity in a complementary strand from another virus, *d* = 3*DL/*4. This yields 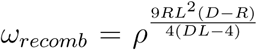. Because mutation and recombination are components of fitness, the probability of each occurring with exclusion of the other at each nucleotide must be accounted. This can be done by replacing *µ* with *Pr*(*µ*\*R*) and *R* with *Pr*(*R*\*µ*). Genome-level fitness for substitution events can be calculated as

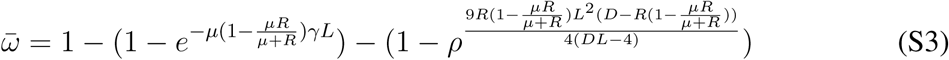

which simplifies to

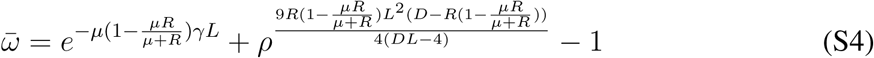

The resulting rate of per nucleotide variability produced is the product of variability accumulation and the average fitness of each genome replicated.

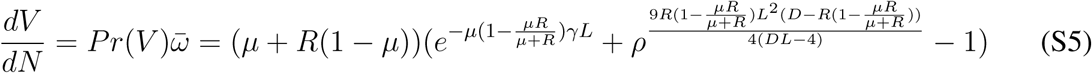

When *µ* and *R* are sufficiently small such that *µR* « *µ, R* (i.e. 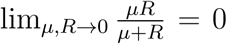), then equation S5 can be approximated as

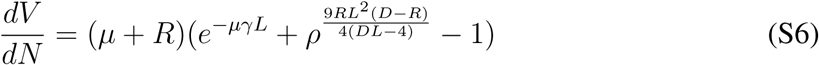

This is given by equation 2 in the main text.

### Parameter values for figure 1

Figure 1 represents a general simulation for ssRNA(+) viruses. To understand the mutation and recombination consequences for this group of viruses, parameter values were approximated from the literature on ssRNA(+) viruses. Because specific estimates for parameter values for each family of viruses within this group is sparse or non-existent, approximate values are given when possible. Qualitative outcomes for each of the panels in figure 1 (main text) are similar across a broad range of possible parameter values, but relationships may differ for other groups of viruses. The parameter values used in the simulation are given in table S1. *r* is the rate constant (*k*_*pol*_) for RNA dependent RNA polymerase in nucleotides per second. This is the speed at which the replicase complex can replicate RNA. *I* is the product of *r* and the amount of additional time (in seconds) required to replicate a strand of RNA outside of the elongation phase plus any additional delay associated with replicating multiple, short strands of RNA rather than fewer, long strands on which multiple replicase complexes could be active simultaneously. The value for *I* chosen was simply an order of magnitude smaller than the large Coronavirus genomes, though this delay has not been quantified in scientific literature. This value is important in determining the genome size at which the maximum rate of production of genetic variability (per second) occurs. *µ* is the mutation rate per nucleotide per replication event by RNA dependent RNA polymerase. This quantity can vary around the value given in table S1, but the value chosen is an appropriate approximation for ssRNA(+) viruses. *R* is the recombination rate per nucleotide per replication event. This value varies considerably in RNA viruses but tends to be high for many ssRNA(+) viruses. *γ* gives the proportion of lethal mutations of the total number of mutations. The vast majority of mutations fall into three categories: neutral, deleterious, or lethal. There is likely variation in the respective proportions of each for viruses, though few attempts have been made to quantify these proportions. In the study referenced, one-third of mutations resulted in viral particles that were not capable of infecting (infectivity assay). This constitutes a lethal mutation and is a good estimate of fitness reduction for cross-species transmission events. *ρ* represents the effective tolerance of proteins to a single recombination event. Misfolding proteins likely constitute lethality or severe reduction in fitness in viruses, so this measure is used to simulate the reduction in fitness in the event of amino acid substitutions due to recombination. *D* is the proportion of dissimilar nucleotides in a co-infecting viral strain or species. Higher dissimilarity results in higher probability of amino acid substitutions due to recombination. 12% dissimilarity is the nucleotide dissimilarity between COVID-19 and SARS-COV.

### Literature review

Peer-reviewed scientific literature was searched for reports on zoonotic viruses transmitted from animals to humans using the Web of Science electronic library database for published reports through 2019. Keywords were used in the search as specified in the main text. Centers for Disease Control and Prevention and World Health Organization reports as well as published compilations of human infectious diseases were also used as references, but reported viruses were cross-validated as zoonotic with other supporting peer-reviewed literature [1, 5, 19]. Zoonotic viruses were identified as having known non-human and human hosts via molecular or serological assay (seroconversion). Viral infections supported by circumstantial evidence sans assay verification were not included. The list of zoonotic viruses was formed after searching for complete genome sequences on the NCBI database [12] then extending to other peer-reviewed full genome sequences if the virus was not found in the NCBI database. This final list included 126 unique viral species. Each viral species was classified according to the Baltimore classification system by mRNA production/genome structure. The compiled list is included as table S3.

### Non-zoonotic virus filtering

The entire NCBI database on viral genomes was downloaded and screened for animal-associated viruses. The category of unidentified viruses was excluded from the screening process. All incomplete sequences were excluded as well. The majority of viruses are labeled with host associations, though in many instances, these labels were found to be inaccurate. Viruses were organized by families and screened initially by known host range. Viral families associated with prokaryotes, archaea, plants, or fungi exclusively were not included. Satellite viruses and viroids were also excluded. Remaining genera and species were manually cross-checked with existing literature to verify the belonging as animal associated. If species were recovered from an animal or had known relatives recovered from an animal, the species was included in the non-zoonotic, animal associated virus list. The animal grouping was assigned due to the difficulty of parsing associations further to vertebrates. Many viruses are recovered from vectors or other invertebrates, yet have known or unknown association with vertebrates. Sometimes viruses have no known association outside of an insect vector, though relatives of the virus are known to infect vertebrate hosts. As an exercise to ensure genome size did not differ substantially between vertebrate and invertebrate-associated viruses, another filtering process was conducted manually whereby each species was cross-validated with existing literature to remove known or likely known to not be associated with a vertebrate host. The total number of non-zoonotic viruses was reduced from 2088 to 1533 in this process. Performing one-tailed Welch’s T-tests with this new data set revealed little departure from previous conclusions. Results are presented in table S2 and Fig. S1.

## Acknowledgments

I would like to thank Kalin Wilson and Paul Bump for providing insightful feedback on this manuscript. I would also like to thank my advisor, Giulio De Leo, for continued support and advice through my PhD.

## Author Contributions

REG contributed to each section of this manuscript.

## Materials and Correspondence

Viral genomes were retrieved from the NCBI viral genome library https://www.ncbi.nlm.nih.gov/genomes/GenomesGroup.cgi. The list of zoonotic virus species with associated genome sizes is provided in the supplement. Python scripts to reproduce the figures is available from the author upon request.

