## Supplemental Material for "Larger viral genome size facilitates emergence of zoonotic diseases"

Richard E. Grewelle,[1,2*]

[1]Department of Biology, Stanford University,

[2]Hopkins Marine Station, Pacific Grove, CA 93950, USA

### Supplemental Text

**Maximizing genetic variability.** The maximum production of genetic variability per unit time is dependent on the virus-specific parameters used in equation 4 (main text). For the parameters used, the maximum can be found under the conditions

$$\frac{d}{dL}\left(\frac{dV}{dt}\right) = 0, \qquad \frac{d^2}{d^2L}\left(\frac{dV}{dt}\right) < 0 \tag{S7}$$

Solving this equation yields a value for the length of the genome (L) near 16000 nt. This value may not reflect the real maximum for ssRNA(+) viruses, as accurate parameter values are difficult to obtain, particularly for $I$. Figure 1 (main text) reflects the qualitative behavior of the curve, which is convex and produces a maximum between 0 and the upper range of genome sizes for RNA viruses. To illustrate, replacing $\mu = 10^{-5}$ with $\mu = 10^{-7}$, reflecting the approximate mutation rate of Coronaviruses, which possess unique proofreading capacity [24], results in a shift from 16000 nt to 22000 nt in the genome size corresponding to the maximum rate of variability production (Figure S2).

**The role of genes and genome size.** Mutation and recombination events are more likely to occur in non-coding regions [17, 32]. Although these regions are not as widespread in viral genomes, large viral genomes are known to contain over 10% non-coding nucleotides. Non-coding elements are not translated into proteins but serve functions such as gene regulation [33, 34]. Intergenic regions are among these non-coding elements and have been identified as hotspots for mutation and recombination, presumably for the weaker selection imposed on these elements compared to genes. Substitution there is less likely to be deleterious or lethal. Large genomes across prokaryotic and eukaryotic taxa are typified by expansion of non-coding, regulatory genetic material, and this is anecdotally true for viruses; larger proportions of non-coding nucleotides are found in viruses with large genomes [34, 35]. Even if the ratio of coding to non-coding nucleotides were fixed as genome size increased, the absolute number of substitution hotspots increases. If the number of genes increases with the size of the genome, the number of intergenic regions increases as well. This says nothing for the size of those intergenic regions, though there are biochemical constraints on the range of sizes these regions can take [36], and it may be reasonably assumed the size of these regions does not scale inversely with the number of them a genome contains. With this premise, the number of nucleotides associated with intergenic regions is positively related to increased genome size, and the number of intergenic regions may well be an indicator of the number of substitution hotspots, and therefore potential for genome-wide substitution. Because the number of intergenic regions is one fewer than the number of genes in a non-segmented genome, the number of genes should recapitulate the likelihood of substitution. In segmented genomes, this relationship may not hold. Figure S3 shows the relationship between genome size and the number of genes, number of intergenic regions, and number of segments in each genome. The number of intergenic regions is calculated as

$$n_{inter} = n_{seg}(\frac{n_{genes}}{n_{seg}} - 1) \tag{S8}$$

where $n_{inter}$, $n_{seg}$, and $n_{genes}$ denote the number of intergenic regions, the number of genomic segments, and the number of genes, respectively. The number of intergenic regions, thus potential for substitutions, is positively related to genome size for groups 1, 2, 4, and 6. Group 1, dsDNA viruses, shows strong correlation between genome size and intergenic regions. This may explain, in part, why differences in genome size between zoonotic and non-zoonotic viruses in group 1 are so great. Although the number of intergenic regions was not related to genome size for group 3, these dsRNA viruses commonly contain segmented genomes, and this segmentation is strongly related to genome size. This relationship demonstrates a fixed segment size near 2500 nucleotides. Generation of variability is likely related to genome size due to the capacity to reassort in dsRNA viruses rather than substitution events through homologous and non-homologous recombination and mutation. Groups 5 and 7 show no relationships between genome size and any of the response variables, suggesting that intergenic regions and reassortment play little role in generation of genetic variability as a function of genome size for these viruses.

**Sampling considerations.** Rigorous screening ensured the inclusion of viruses that have known animal hosts and have elicited immune response in humans. Viruses for which there is extensive debate about the host were excluded. An example is human picobirnavirus. Although previously identified in stool samples in rats and humans, this virus shares sequence homology with prokaryotic viruses and is expected to be a prokaryotic virus [37]. Further work is needed to determine whether this virus is truly zoonotic or exists in hosts with shared enteric flora. For viruses identified as zoonotic, sequences were retrieved from [12], and if the sequences were unavailable, complete sequence length was found from other published sources. All sequences in [12] corresponding to known zoonoses were used in the zoonotic group for analysis. This includes one isolate (England Coronavirus) that may not be considered a separate species. Species-level identification is notoriously difficult for viruses due to rapid mutation and recombination rates, and this likely has resulted in inclusion of instances of species redundancy in the non-zoonotic data set, though these instances are perceived to be rare. The NCBI Viral Genome library is filtered and curated to eliminate redundancies in species identification.

The viral sequences in [12] encompass a vast array of viruses across host taxa, and considerable effort was given to filter non-animal viruses. All viruses of unknown classification were removed. Viral families tend to cluster with host phyla, so proper classification by viral family was possible in the vast majority of cases. The vast majority of viruses have yet to be detected, and the existing viral sequences are certainly biased toward those associated with human and domestic animal and plant pathology. Viruses were grouped by the Baltimore classification system, which is based on the method of mRNA synthesis and the structure of the viral genome. This level of grouping was chosen over higher level grouping (DNA vs RNA) because there are distinct differences in genome size that occur due to method of mRNA synthesis, and the constraints replication and mRNA synthesis impose on genome structure are best conserved within categories defined by the Baltimore classification system. Lower level grouping was avoided because of the difficulty of taxonomic classification of viruses at lower levels. Additionally, as is evident by the list of zoonotic viruses, some clades of viruses are more likely to exhibit broad host ranges, and parsing genome size effects on the likelihood of cross-species transmission is less reliable within clades than between clades with the available sample sizes.

### Supplemental Figures

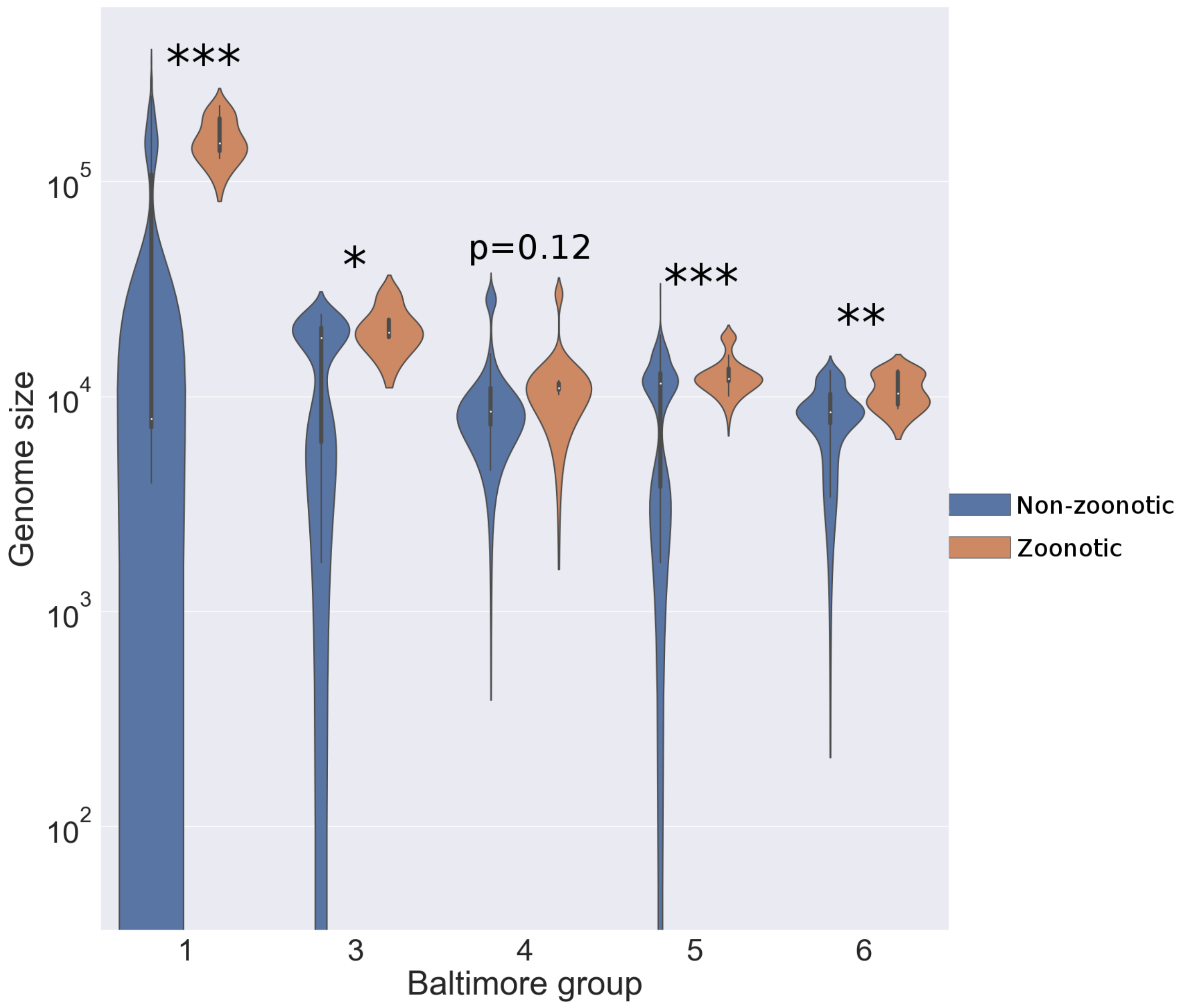


Figure S1: Violin plots comparing genome sizes between non-zoonotic vertebrate-associated viruses and zoonotic viruses. Zoonotic virus genome sizes inhabit the upper ranges of values in each group.

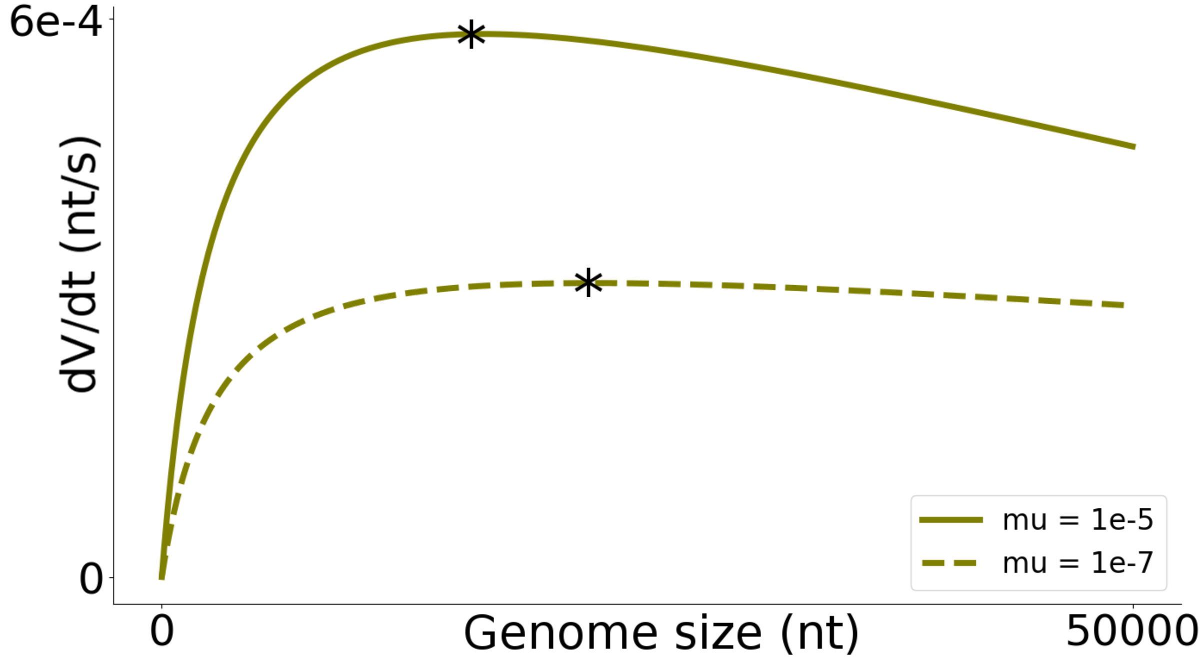


Figure S2: Lower rates of mutation favor larger genome sizes for maximum rates of production of variability per unit time per infecting virion. Asterisks indicate maxima for each curve.

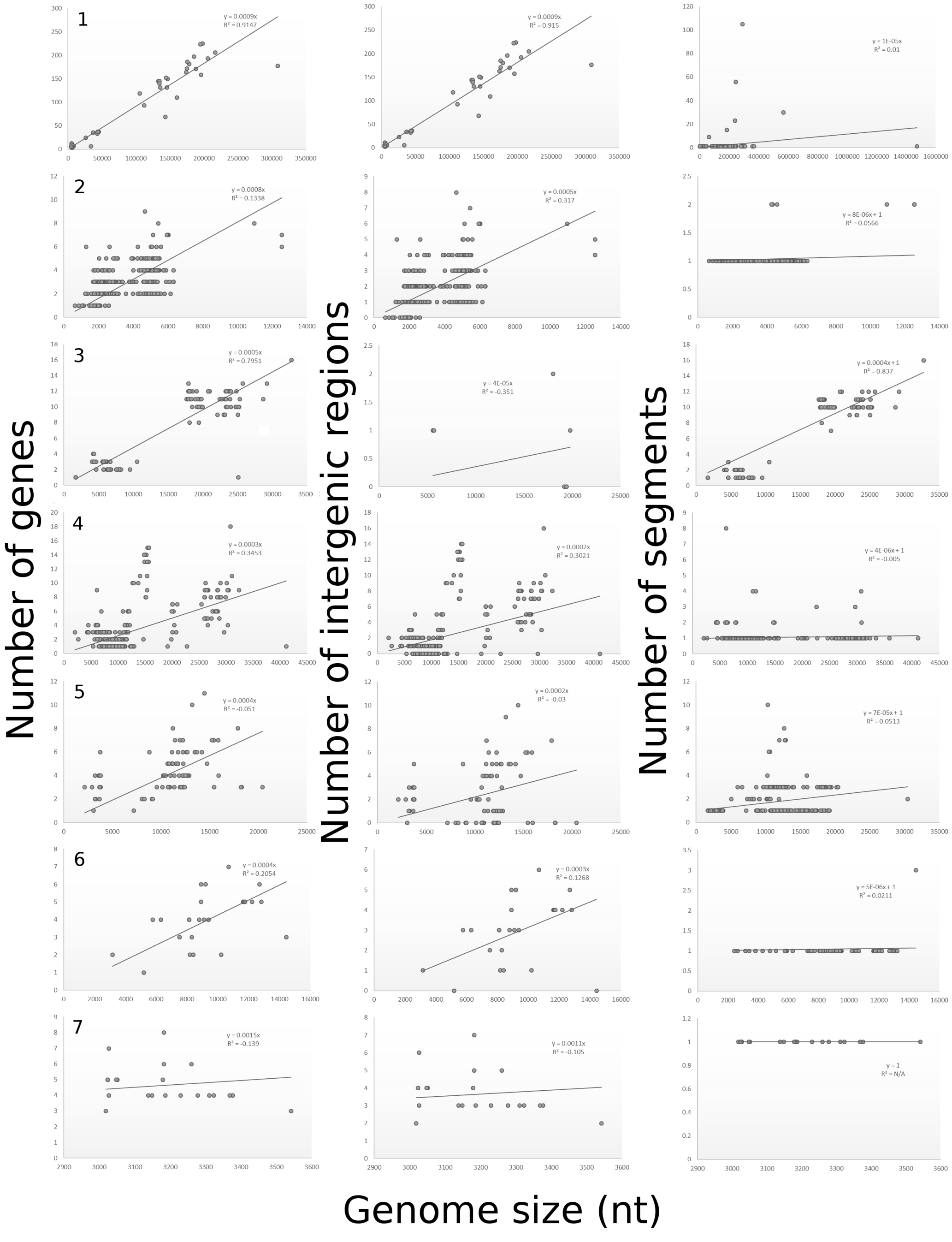


Figure S3: Scaling of number of genes, intergenic regions, and genomic segments with genome size. 7 Baltimore groups are presented, one for each row: dsDNA (1), ssDNA (2), dsRNA (3), ssRNA+ (4), ssRNA- (5), ssRNA-RT (6), dsDNA-RT (7). Intercepts of linear regressions are fixed at the origin (or at y=1 for the last column). Negative $R^2$ values reveal a correlation between the regression line and the data that is weaker than a line with zero slope and fitted intercept.

### Supplemental Tables

| Parameter | Description | Value | Reference |
|---|---|---|---|
| r | RNA polymerization rate | $40\ nt\ s^{-1}$ | [38] |
| I | Initiation time (nt equiv.) | $3000\ nt$ | NA |
| $\mu$ | Mutation rate | $10^{-5}\ nt^{-1}$ | [14] |
| R | Recombination rate | $10^{-5}\ nt^{-1}$ | [27] |
| $\gamma$ | Proportion of lethal mutations | 0.33 | [18] |
| $\rho$ | Tolerance to recombination | 0.79 | [17] |
| D | Genetic dissimilarity | 0.12 | [25] |

Table S1: Approximate values for parameters used in figure 1 (main text).

| Group | Genome | Vertebrate Virus Genome Size | Zoonotic Virus Genome Size | P-value |
|---|---|---|---|---|
| 1 | dsDNA | 54123 ± 3280 (n=540) | 165175 ± 12595 (n=8) | $<$0.0001 |
| 3 | dsRNA | 15495 ± 970 (n=60) | 21828 ± 2493 (n=4) | 0.05 |
| 4 | ssRNA(+) | 11397 ± 327 (n=468) | 12575 ± 920 (n=41) | 0.12 |
| 5 | ssRNA(-) | 10225 ± 253 (n=393) | 12964 ± 336 (n=64) | $<$ 0.0001 |
| 6 | ssRNA-RT | 8600 ± 308 (n=72) | 10949 ± 637 (n=9) | 0.003 |

Table S2: Comparison of viral genome size between known zoonotic viruses and all other vertebrate viruses. Comparisons are made within Baltimore groups classified according to genomic structure. Mean and standard errors for each group are reported. Little change is observed between this table and table 1 in the main text.

Table S3: Zoonotic Viruses

| Viral Genus | Species Common Name | Viral Group | Genome Size (ref.) |
|---|---|---|---|
| Orthopoxvirus | Buffalopox virus | 1 | 195630 [39] |
| Orthopoxvirus | Monkeypox virus | 1 | 196858 [40] |
| Parapoxvirus | Bovine papular stomatitis | 1 | 134431 [41] |
| Parapoxvirus | Cowpox | 1 | 224499 [42] |
| Parapoxvirus | Orf virus | 1 | 139962 [42] |
| Parapoxvirus | Pseudocowpox virus | 1 | 145289 [43] |
| Parapoxvirus | Sealpox | 1 | 127941 [44] |
| Simplexvirus | Herpes virus B | 1 | 156789 [45] |
| Coltivirus | Colorado tick fever | 3 | 29174 [46] |
| Orbivirus | Orungo virus | 3 | 18894 [47] |
| Rotavirus | Rotavirus A | 3 | 18562 [48] |
| Seadornavirus | Banna | 3 | 20682 [49] |
| Alphavirus | Barmah Forest virus | 4 | 11488 [50] |
| Alphavirus | Chikungunya | 4 | 11826 [51] |
| Alphavirus | Eastern equine encephalitis | 4 | 11675 [52] |
| Alphavirus | Everglades virus | 4 | 11395 [53] |
| Alphavirus | Getah virus | 4 | 11597 [12] |
| Alphavirus | Mayaro virus | 4 | 11411 [12] |
| Alphavirus | O’nyong-nyong virus | 4 | 11835 [54] |
| Alphavirus | Ross River virus | 4 | 11657 [55] |
| Alphavirus | Semliki Forest virus | 4 | 11442 [56] |
| Alphavirus | Sindbis virus | 4 | 11703 [57] |
| Alphavirus | Venezuelan equine encephalitis virus | 4 | 11444 [58] |
| Alphavirus | Western equine encephalitis virus | 4 | 11484 [59] |
| Aphthovirus | Foot and mouth disease | 4 | 8201 [60] |
| Betacoronavirus | England isolate, MERS Coronavirus | 4 | 30111 [12] |
| Betacoronavirus | MERS Coronavirus | 4 | 30119 [61] |
| Betacoronavirus | Novel Coronavirus | 4 | 29882 [62] |
| Betacoronavirus | SARS Coronavirus | 4 | 29751 [63] |
| Cardiovirus | Encephalomyocarditis virus | 4 | 7835 [64] |
| Flavivirus | Alkhurma virus | 4 | 10775 [65] |
| Flavivirus | Dengue fever 1 | 4 | 10735 [66] |
| Flavivirus | Dengue fever 2 | 4 | 10723 [67] |
| Flavivirus | Dengue fever 3 | 4 | 10707 [68] |
| Flavivirus | Dengue fever 4 | 4 | 10649 [69] |
| Flavivirus | Edge Hill virus | 4 | 10206 [70] |

Table S3 (cont.): Zoonotic Viruses

| **Viral Genus** | **Species Common Name** | **Viral Group** | **Genome Size (ref.)** |
|---|---|---|---|
| Flavivirus | European tick-borne encephalitis | 4 | 11141 [71] |
| Flavivirus | Far eastern tick-borne encephalitis | 4 | 10471 [72] |
| Flavivirus | Ilheus virus | 4 | 10755 [73] |
| Flavivirus | Japanese encephalitis virus | 4 | 10976 [74] |
| Flavivirus | Kunjin virus | 4 | 10644 [75] |
| Flavivirus | Kyasanur Forest disease virus | 4 | 10774 [76] |
| Flavivirus | Murray Valley encephalitis virus | 4 | 11014 [77] |
| Flavivirus | Omsk virus | 4 | 10787 [78] |
| Flavivirus | Powassan virus | 4 | 10839 [79] |
| Flavivirus | St. Louis encephalitis virus | 4 | 10940 [80] |
| Flavivirus | Usutu virus | 4 | 11066 [81] |
| Flavivirus | Wesselsbron virus | 4 | 10814 [82] |
| Flavivirus | West Nile Virus | 4 | 10962 [83] |
| Flavivirus | Yellow fever virus | 4 | 10862 [84] |
| Flavivirus | Zika virus | 4 | 10807 [85] |
| Orthohepevirus | Hepatitis E | 4 | 7176 [86] |
| Parechovirus | Ljungan virus | 4 | 7590 [87] |
| Alphainfluenzavirus | Influenza A virus (2009(H1N1)) | 5 | 13158 [88] |
| Alphainfluenzavirus | Influenza A virus (1996(H5N1)) | 5 | 13590 [89] |
| Alphainfluenzavirus | Influenza A virus (1999(H9N2)) | 5 | 13498 [90] |
| Alphainfluenzavirus | Influenza A virus (1968(H2N2)) | 5 | 13460 [91] |
| Alphainfluenzavirus | Influenza A virus (2004(H3N2)) | 5 | 13627 [91] |
| Alphainfluenzavirus | Influenza A virus (1934(H1N1)) | 5 | 13588 [92] |
| Alphainfluenzavirus | Influenza A virus (2013(H7N9)) | 5 | 13191 [93] |
| Orthoavulavirus | Newcastle disease | 5 | 15192 [94] |
| Orthobornavirus | Borna disease 1 | 5 | 8910 [95] |
| Orthobornavirus | Borna disease 2 | 5 | 8908 [96] |
| Ebolavirus | Bundibugyo Ebola virus | 5 | 18940 [12] |
| Ebolavirus | Reston Ebola virus | 5 | 18891 [97] |
| Ebolavirus | Sudan Ebola virus | 5 | 18875 [98] |
| Ebolavirus | Tai Forest Ebola virus | 5 | 18935 [99] |
| Ebolavirus | Zaire Ebola virus | 5 | 18959 [100] |
| Henipavirus | Hendra virus | 5 | 18234 [101] |
| Henipavirus | Nipah virus | 5 | 18246 [102] |
| Lyssavirus | Australian bat lyssavirus | 5 | 11822 [103] |
| Lyssavirus | Duvenhage virus | 5 | 11976 [104] |

Table S3 (cont.): Zoonotic Viruses

| Viral Genus | Species Common Name | Viral Group | Genome Size (ref.) |
|---|---|---|---|
| Lyssavirus | European bat lyssavirus type 1 | 5 | 11966 [105] |
| Lyssavirus | European bat lyssavirus type 2 | 5 | 11930 [105] |
| Lyssavirus | Mokola virus | 5 | 11940 [106] |
| Lyssavirus | Rabies virus | 5 | 11932 [107] |
| Mammarenavirus | Chapare virus | 5 | 10464 [108] |
| Mammarenavirus | Guanarito | 5 | 10424 [12] |
| Mammarenavirus | Junin virus | 5 | 10525 [12] |
| Mammarenavirus | Lassa virus | 5 | 10697 [109] |
| Mammarenavirus | Lujo virus | 5 | 10352 [110] |
| Mammarenavirus | Lymphocytic choriomeningitis virus | 5 | 10056 [111] |
| Mammarenavirus | Machupo virus | 5 | 10635 [12] |
| Mammarenavirus | Sabia virus Brazilian hemorrhagic fever | 5 | 10499 [12] |
| Mammarenavirus | Whitewater Arroyo virus | 5 | 10448 [112] |
| Marburgvirus | Marburg virus | 5 | 19114 [113] |
| Orthobunyavirus | Cache Valley virus | 5 | 12283 [12] |
| Orthobunyavirus | California encephalitis | 5 | 12466 [114] |
| Orthobunyavirus | Guama | 5 | 12123 [115] |
| Orthobunyavirus | Guaroa virus | 5 | 12265 [116] |
| Orthobunyavirus | Jamestown Canyon virus | 5 | 12461 [114] |
| Orthobunyavirus | Kairi virus | 5 | 12497 [12] |
| Orthobunyavirus | LaCrosse virus | 5 | 12490 [12] |
| Orthobunyavirus | Oropouche virus | 5 | 11985 [12] |
| Orthobunyavirus | Tahyna virus | 5 | 12446 [117] |
| Orthohantavirus | Andes virus | 5 | 11909 [118] |
| Orthohantavirus | Bayou hantavirus | 5 | 12189 [12] |
| Orthohantavirus | Dobrava virus | 5 | 11840 [119] |
| Orthohantavirus | Hantaan | 5 | 11845 [120] |
| Orthohantavirus | Monongahela virus | 5 | 12314 [121] |
| Orthohantavirus | Muju virus | 5 | 12027 [122] |
| Orthohantavirus | Prospect Hill orthohantavirus | 5 | 11941 [123] |
| Orthohantavirus | Puumala virus | 5 | 12062 [12] |
| Orthohantavirus | Seoul virus | 5 | 11950 [12] |
| Orthohantavirus | Sin Nombre virus | 5 | 12317 [124] |
| Orthohantavirus | Tula virus | 5 | 12066 [125] |
| Orthonairovirus | Crimean-Congo hemorrhagic fever | 5 | 19146 [126] |
| Phlebovirus | Bhanja virus | 5 | 11511 [127] |

Table S3 (cont.): Zoonotic Viruses

| Viral Genus | Species Common Name | Viral Group | Genome Size (ref.) |
|---|---|---|---|
| Phlebovirus | Rift Valley fever virus | 5 | 11979 [128] |
| Phlebovirus | Toscana (sandfly) virus | 5 | 12488 [12] |
| Phlebovirus | Turkey (sandfly) virus | 5 | 12603 [129] |
| Rubulavirus | Menangle virus | 5 | 15516 [130] |
| Rubulavirus | Tioman virus | 5 | 15522 [131] |
| Vesiculovirus | Chandipura virus | 5 | 11120 [132] |
| Vesiculovirus | Vesicular stomatitis infection (Alagoas) | 5 | 11070 [133] |
| Vesiculovirus | Vesicular stomatitis infection (IN) | 5 | 11162 [134] |
| Vesiculovirus | Vesicular stomatitis infection (NJ) | 5 | 11123 [135] |
| Deltaretrovirus | Primate T-lymphotropic virus 1 | 6 | 9028 [136] |
| Deltaretrovirus | Primate T-lymphotropic virus 4 | 6 | 8791 [12] |
| Lentivirus | Human immunodeficiency virus 1 | 6 | 9181 [137] |
| Lentivirus | Human immunodeficiency virus 2 | 6 | 10359 [138] |
| Lentivirus | Simian immunodeficiency virus | 6 | 9623 [139] |
| Spumavirus | Simian Foamy virus | 6 | 13246 [140] |
| Spumavirus | Guenon simian foamy virus | 6 | 13072 [141] |
| Spumavirus | Gorilla simian foamy virus | 6 | 12258 [142] |
| Spumavirus | Rhesus macaque foamy virus | 6 | 12983 [12] |

32. Men, R., Bray, M., Clark, D., Chanock, R. M. & Lai, C.-J. Dengue type 4 virus mutants containing deletions in the 3'noncoding region of the RNA genome: analysis of growth restriction in cell culture and altered viremia pattern and immunogenicity in rhesus monkeys. *Journal of virology* **70,** 3930–3937 (1996).

33. Tycowski, K. T. *et al.* Viral noncoding RNAs: more surprises. *Genes & development* **29,** 567–584 (2015).

34. Mahmoudabadi, G. & Phillips, R. A comprehensive and quantitative exploration of thousands of viral genomes. *Elife* **7,** e31955 (2018).
